# GATA4 and GATA6 loss-of-expression is associated with extinction of the classical programme and poor outcome in pancreatic ductal adenocarcinoma

**DOI:** 10.1101/2021.08.09.455642

**Authors:** Mónica P. de Andrés, Richard Jackson, Christian Pilarsky, Anna Melissa Schlitter, Eithne Costello, William Greenhalf, Paula Ghaneh, Thomas Knösel, Daniel Palmer, Petra Rümmele, Wilko Weichert, Markus Büchler, Thilo Hackert, John P. Neoptolemos, Núria Malats, Paola Martinelli, Francisco X. Real

**Affiliations:** Epithelial Carcinogenesis Group, Spanish National Cancer Research Centre-CNIO, Madrid, Spain; Institute of Translational Medicine, University of Liverpool, Liverpool, United Kingdom; Department of Surgery, Universitätsklinikum Erlangen, Erlangen, Germany; Institute of Pathology, School of Medicine, Technische Universität München, Munich, Germany; German Cancer Consortium (DKTK), German Cancer Research Center (DKFZ), Heidelberg, Germany; Department of Molecular and Clinical Cancer Medicine, University of Liverpool; Institute of Pathology, Ludwig-Maximilians-University Munich, Munich, Germany; Pathologisches Institut, University of Erlangen, Erlangen, Germany; Institute of Pathology of the Technical University Munich, Munich, Germany; Department of General, Visceral, and Transplantation Surgery, University of Heidelberg, Heidelberg, Germany; Genetic and Molecular Epidemiology Group, Spanish National Cancer Research Centre-CNIO, Madrid, Spain; CIBERONC, Madrid, Spain; Institute of Cancer Research, Clinic for Internal Medicine I, Medical University of Vienna, Austria; Departament de Ciències Experimentals i de la Salut Universitat Pompeu Fabra, Barcelona, Spain

**Author notes:** **Corresponding author:** Francisco X. Real.

## Abstract

**Objective:** GATA6 is a master regulator of pancreatic differentiation and a key regulator of the classical phenotype in pancreatic ductal adenocarcinoma (PDAC). Low GATA6 expression is associated with poor patient outcome. *GATA4* is the second most expressed GATA factor in the pancreas. The aim was to assess whether, and how, GATA4 contributes to PDAC phenotype and to analyze the association of expression with clinical outcome.

**Design:** We analyzed PDAC transcriptomic data, stratifying cases according to *GATA4* and *GATA6* expression, and identified differentially expressed genes and pathways. A multicenter TMA study to assess GATA4 and GATA6 expression in PDAC samples (n=745) from patients undergoing tumour resection was performed using immunohistochemistry with antibodies of validated specificity. GATA4 and GATA6 levels were dichotomized into high/low categorical variables; association with outcome was assessed using univariable and multivariable Cox regression models.

**Results:** Subtype classification using transcriptomic data revealed that *GATA4* mRNA is enriched in classical, compared to basal-like tumours. We classified samples in 4 groups as high/low for *GATA4* and *GATA6*. Reduced expression of *GATA4* did not have a major transcriptional impact. However, concomitant low expression of *GATA4* enhanced the transcriptomic effects of *GATA6* low expression. Reduced expression of both proteins in tumours was associated with the worst patient survival. *GATA4* and *GATA6* expression significantly decreased in metastases and negatively correlated with basal markers.

**Conclusions:** Our analyses uncover a cooperative interaction between *GATA4* and *GATA6* to maintain the classical PDAC phenotype and provide compelling clinical rationale for assessing their expression as biomarkers of poor prognosis.

**SUMMARY BOX:** *What is already known about this subject?:*

- Patients with classical-type PDAC have a better outcome
- Retrospective analyses suggest that classical-type PDAC is more sensitive to 5-FU-based chemotherapy
- GATA6 is a surrogate biomarker of classical tumours and its expression is associated with better survival
- GATA4 and GATA6 have overlapping and unique functions during pancreatic and gastrointestinal development

*What are the new findings?:*

- Tumours displaying only low GATA4 expression have a transcriptomic profile similar to those with preserved expression of both transcription factors
- Combined low expression of GATA4 and GATA6 has the highest transcriptomic impact
- In a large multicenter tissue microarray study, patients with tumours showing low expression of both GATA4 and GATA6 have the worst overall survival
- Low expression of GATA4 and GATA6 is an independent predictor of survival in patients with resectable PDAC
- GATA4 levels are down-regulated in liver metastases and are negatively correlated with basal markers such as KRT5/6, KRT14, and TP63

*How might it impact on clinical practice in the foreseeable future?:*

- The combined assessment of GATA4 and GATA6 expression may improve prognostic stratification of patients with PDAC
- Prospective studies should confirm whether GATA4 and GATA6 expression is also predictive of response to chemotherapy in PDAC patients

## INTRODUCTION

Pancreatic ductal adenocarcinoma (PDAC) remains a deadly malignancy with a 5-year survival rate of only 10%[1]. Non-specific symptoms, late diagnosis, and aggressive biology, among other factors, account for the poor outcome[2]. Only 20% of patients have resectable tumours at presentation and the 5-year-survival rate after surgery alone with radical intent is only 8%, rising to 30-50% using adjuvant combination chemotherapy after resection[3,4]. The standard-of-care for patients with resectable disease is adjuvant therapy with modified 5-fluorouracil, leucovorin, irinotecan and oxaliplatin (FOLFIRINOX) or gemcitabine with capecitabine[4,5]. FOLFIRINOX or gemcitabine (with or without Abraxane) are used in locally advanced and metastatic disease[6,7]. Neoadjuvant chemotherapy is increasingly used in patients with borderline resectable PDAC and it may impact on patient outcome since PDAC is considered to be a systemic disease[8]. An improved understanding of the biological basis of PDAC aggressiveness is needed.

Tumour stratification through genomic analysis has unraveled a quite homogenous landscape, where most tumours share recurrent alterations in four genes (*KRAS, TP53, CDKN2A, SMAD4*). In this background, each tumour has a private constellation of additional genomic alterations, rendering PDAC a highly heterogeneous disease. Transcriptomic profiling of PDAC samples has allowed a molecular classification consistently identifying two main tumour categories: classical and basal[9–12]. High expression of adhesion-and epithelial-associated genes[9], up-regulation of GATA6 and HNF1A, and better prognosis are hallmarks of the classical subtype[9,10] (also designated as “pancreatic progenitor”[11], and “classical A/B”[12]). Basal tumours (“quasimesenchymal”[9], “basal-like”[10], “squamous”[11], and “basal-like A/B”[12]) are characterized by down-regulation of endodermal transcription factors (e.g. GATA6, HNF1A, and PDX1) and epithelial markers (e.g.CDH1), poor survival, high expression of keratins of stratified epithelium (e.g. KRT14 and KRT5/6)[10], and of ΔNP63[11]. Increasing evidence suggests that PDAC subtypes are continuous rather than dichotomous, resulting both from the co-occurrence of cells with classical and basal features in the same tumour as well as the existence of a continuum of phenotypes at the single cell level[3,12,13]. GATA6 has emerged as a master regulator of canonical differentiation programmes both in normal pancreas and in PDAC[3,14]. A *GATA6* superenhancer is involved in the classical PDAC phenotype[15] and, among all up-regulated pancreatic lineage transcription factors in this subtype, *GATA6* has been found to be recurrently amplified[12]. GATA6 expression is a surrogate biomarker for classical PDAC[16] and is associated with response to adjuvant therapy[3,16].

The GATA family of transcription factors comprises six zinc finger proteins that bind the consensus DNA sequence (T/A)GATA(A/G). They play complex roles in embryonic development and cancer[17]. GATA6 and GATA4 are key regulators of heart development: *GATA6* haploinsufficiency causes congenital heart defects in humans[18,19] and *GATA4* mutations cause cardiac septal defects and dilated cardiomyopathy[20,21]. Regarding the pancreas, *GATA6* mutations can lead to a broad phenotypic spectrum, ranging from adult-onset diabetes to pancreatic hypoplasia/agenesis[18,19]. *GATA4* inactivating mutations and deletions cause neonatal or childhood-onset diabetes, with variable degrees of exocrine insufficiency[22]. *Gata4* and *Gata6* also share partially redundant functions during pancreas organogenesis, since concomitant inactivation results in pancreas agenesis while inactivation of only one of them does not impede pancreas formation[23,24]. *Gata4* inactivation in the Pdx1-expressing domain during mouse development results in pancreatic heterotopia in the stomach[25]. Accordingly, GATA4 and GATA6 control shared developmental programmes but also have gene-specific functions.

Here, we show that GATA4 is expressed at high levels in normal human adult pancreas and a subset of PDAC. Through bioinformatics analysis of available transcriptomic datasets of PDAC[11,12], we find that GATA4 mRNA is down-regulated in the basal subtype and expression of both GATA transcripts is positively correlated. At the molecular level, low expression of GATA4 amplifies the transcriptomic effect of GATA6 down-regulation. Orthogonal validation of GATA4 and GATA6 protein levels in a large series of clinically-annotated PDAC samples from patients undergoing surgical resection with curative intent in the context of clinical trials or standard of care treatment, revealed that concomitant low expression of GATA6 and GATA4 is consistently associated with worse patient outcome than GATA6 low expression alone. Finally, we show that low GATA4 and GATA6 expression is associated with liver metastasis. Our studies provide novel clinical insights supporting a cooperative interaction between GATA4 and GATA6 to sustain the classical PDAC phenotype and uncover GATA4 as a novel marker of tumour progression that may contribute to refine patient stratification.

## METHODS AND MATERIALS

### Details are provided as Supplementary Information

The following PDAC RNA-Seq datasets were used in the analyses: ICGC (n=96)[11] and PanCuRx (n=247).

### Tumour classification

To establish tumour categories, the distribution of GATA4 and GATA6 expression values was assessed. Four categories were defined in both datasets: GATA4 high/GATA6 high (G4^Hi^/G6^Hi^), GATA4 low/GATA6 high (G4^Lo^/G6^Hi^), GATA4 high/GATA6 low (G4^Hi^/G6^Lo^), and GATA4 low/GATA6 low (G4^Lo^/G6^Lo^).

### Differential gene expression analysis, Weighted Gene Correlation Network Analysis (WGCNA), and Gene Set Enrichment Analysis (GSEA)

Differential gene expression analysis was performed using Comparative Marker Selection (version 10.1) and Comparative Marker Selection Viewer (version 9) tools[26] through GenePattern using the default parameters. The R package WGCNA was applied to analyze the PanCuRx dataset. The topological overlap matrix (TOM) algorithm was used to identify modules of densely interconnected genes[27]. GSEA was performed using the tool “Investigate Gene Sets” provided by the Molecular Signatures Database v7.4[28].

### Patient cohorts and clinical information

Tissue samples for tissue micro arrays (TMAs) were obtained from patients who underwent pancreatic resection for PDAC. All study sites provided tissue microarrays (TMAs) containing tumour samples from patients who underwent surgical resection. A summary of relevant information is provided in Supplementary Tables 1 and 2. Four cohorts were included from Charité University Hospital[29], Klinikum rechts der Isar, TU München, Germany[30], Regensburg and Jena Institutes of Pathology [31], and the ESPAC-3 trial[32]. Ethics Committee approval information is provided in the Supplementary Material.

### Survival analyses

The primary outcome was overall survival (OS); estimates of OS were obtained using Kaplan Meier curves. Univariable and multivariable analyses were performed.

## RESULTS

### GATA4 and GATA6 expression is associated with the classical phenotype in PDAC

To assess the expression of GATA transcripts in normal pancreas, we examined the transcriptomes of 228 human pancreas samples from the Genotype-Tissue Expression (GTEx) Project[33]. GATA4 and GATA6 are the two main GATA factors expressed in the normal pancreas (Figure 1A). Immunohistochemical analysis of normal human pancreas using specific antibodies revealed that GATA4 is detected only in acinar cells whereas GATA6 is detected in acinar, ductal and islet cells (Figure 1B).

**Figure 1.**
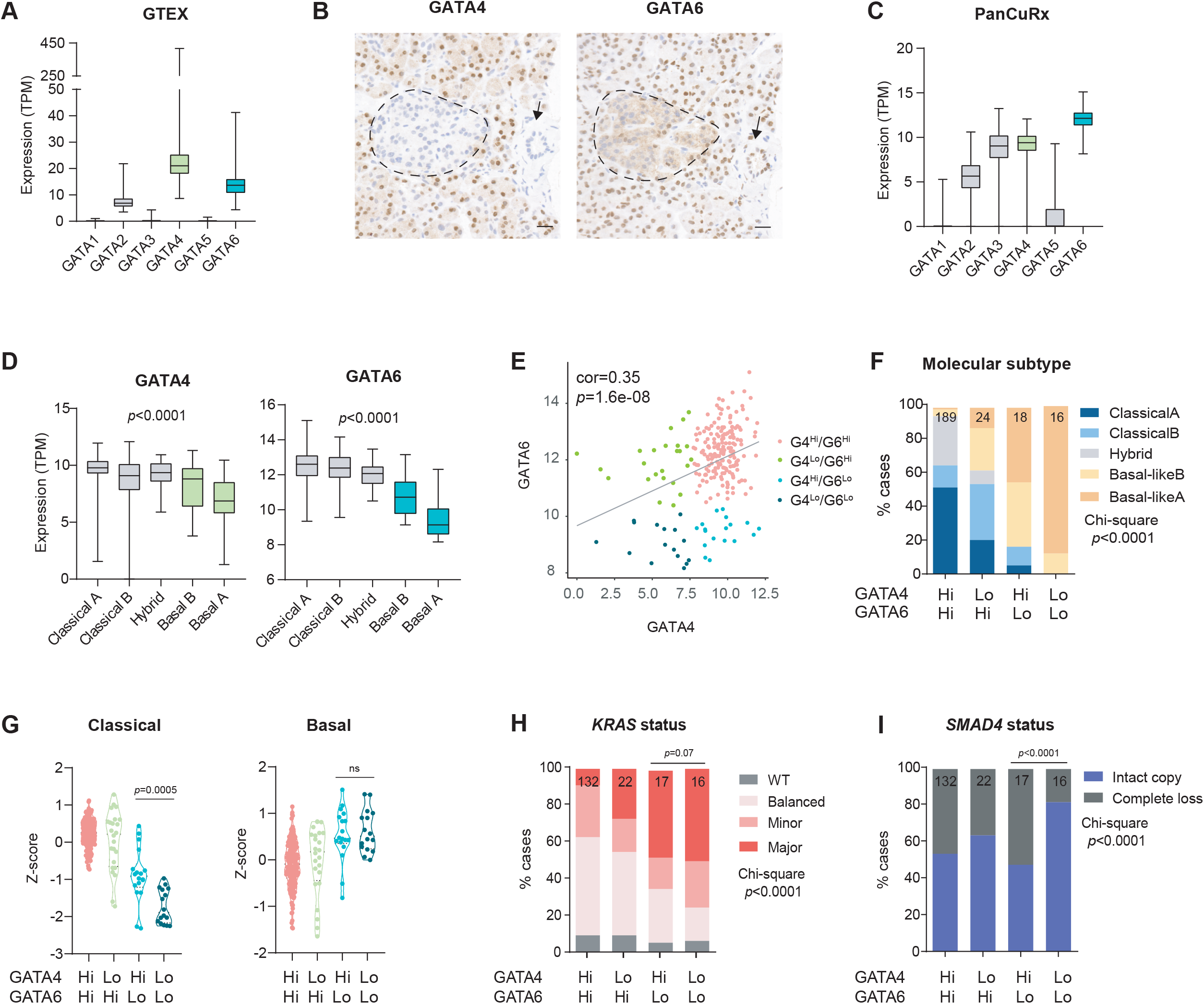
GATA4 and GATA6 are expressed in pancreatic cancer and are down-regulated in the basal subtype. (A) mRNA expression of GATA1-6 in normal human pancreas. Data was extracted from GTEx (see methods). (B) Representative IHC images of the expression of GATA4 and GATA6 in normal human pancreas. GATA6 is expressed in acinar, islet (dashed encircled area) and ductal cells (black arrow) whereas GATA4 is only detected in acinar cells. Scale bar: 20uM. (C) mRNA expression of GATA1-6 in the PanCuRx PDAC dataset. (D) Gene expression levels of GATA4 and GATA6 among PDAC subtypes from the PanCuRx dataset (boxplots are annotated by a Kruskal-Wallis P value). (E) Scatter plot showing a positive correlation between GATA4 and GATA6 mRNA expression (Pearson correlation) and sample classification after GATA4 and GATA6 data dichotomization in the PanCuRx dataset (see methods). (F) Stacked bar plot representing PDAC subtype distribution according to GATA4/GATA6 expression categories (Chi-square test, *P<*0.0001). (G) Z-score of the “Classical” and “Basal” signatures[10] for each of the samples according to the GATA4/GATA6 expression categories; Z-score was calculated as the mean expression (adjusted by Z-score) of the genes in the signature (Kruskal-Wallis *P<*0.0001, two-sided Mann–Whitney test G4^Hi^/G6^Lo^ vs G4^Lo^/G6^Lo^). (H, I) Stacked bar plot representing *KRAS* status or *SMAD4* status according to the GATA4/GATA6 expression categories in the PanCuRx samples (Chi-square test, *P<*0.0001).

To compare GATA4 and GATA6 expression in pancreatic cancer, we took advantage of the transcriptome data from the PanCuRx study that used laser-microdissection to enrich epithelial cells from primary and metastatic tumours[12]. GATA4 and GATA6 are also the main GATA genes expressed in this PDAC series and their levels show a moderate positive correlation (r=0.35, p-value = 1.617e-08; Pearson’s test) (Figure 1C,E). We reproduced these findings using transcriptomic data from the Australian ICGC study of non-microdissected PDAC samples[11] (r=0.39, p-value=8.829e-05) (Supplementary Figure 1A,B). Classification of tumour samples according to molecular subtypes revealed that basal tumours display significantly lower levels of GATA4 and GATA6 mRNAs than classical tumours (Figure 1D).

Next, we classified samples from the PanCuRx study according to GATA4 and GATA6 expression levels; thresholds were established according to the distribution of the data (Supplementary Figure 1C, see Methods). Tumours were classified in four groups resulting from dichotomizing each covariate, as follows: GATA4 high/GATA6 high (G4^Hi^/G6^Hi^) (n=189), GATA4 low/GATA6 high (G4^Lo^/G6^Hi^) (n=24), GATA4 high/ GATA6 low (G4^Hi^/G6^Lo^) (n=18), and GATA4 low/GATA6 low (G4^Lo^/G6^Lo^) (n=16) (Figure 1E, Supplementary Figure 1D).

Using the molecular classification proposed in the PanCuRx study, we examined the association of GATA gene expression with tumour subtypes (classical, hybrid, or basal). The G4^Hi^/G6^Hi^ group was highly enriched in the Classical or Hybrid subtypes (35% and 29%, respectively). Tumours showing selective down-regulation of GATA4 mRNA were also highly enriched in these subtypes, whereas those showing selective down-regulation of GATA6 mRNA were highly enriched in the Basal subtypes (83%). All tumours showing low expression of both GATA6 and GATA4 belonged to the Basal subtype and 88% of them were classified as Basal-like A (Figure 1F). To refine these analyses, we examined the expression of classical and basal genes in the epithelial compartment using the signature reported by Moffit et al[10]: selective reduction of GATA6, but not of GATA4, was associated with a significantly lower Z-score of the classical signature. Tumours showing concomitant low GATA4 and GATA6 expression showed the lowest classical score. In contrast, the Z-score of the Basal gene signature was not significantly different between the G4^Hi^/G6^Lo^ and the G4^Lo^/G6^Lo^ groups (Figure 1G).

Next, we analyzed the status or *KRAS* and *SMAD4* in the four groups using whole-genome sequencing (WGS) data from the PanCuRx study. Tumours with homologous recombination defects and DNA mismatch repair defects [24%] were excluded due to unique mutational signatures[12]. The prevalence of *KRAS*-WT tumours was similar across groups but low expression of GATA4 and/or GATA6 was associated with increased imbalance of the mutant vs wild type allele (Chi-square test, *P<*0.0001; G4^Hi^/G6^Lo^ vs G4^Lo^/G6^Lo^, two-sided Fisher’s exact test *P=*0.076) (Figure 1H). Interestingly, 80% of the G4^Lo^/G6^Lo^ tumours had an intact copy of *SMAD4* (Chi-square test, *P<*0.0001; G4^Hi^/G6^Lo^ vs G4^Lo^/G6^Lo^, two-sided Fisher’s exact test *P<*0.0001) (Figure 1I).

In sum, GATA4 is highly expressed in normal human pancreas and in human PDAC, where it is down-regulated in basal tumours. Low expression of GATA4 on its own does not have a major impact on basal genes, suggesting that it participates in the maintenance of the canonical pancreatic differentiation programme when GATA6 expression is lost.

### Low GATA4 expression is associated with enhanced transcriptomic effects of GATA6 down-regulation

To identify the mechanisms that might explain the differential contribution of GATA4 and GATA6 to the maintenance of the classical phenotype, we first performed a differential gene expression analysis using genome wide RNA-Seq data from the PanCuRx study. Taking the G4^Hi^/G6^Hi^ group as reference, 318 genes (87 up-and 231 down-) were significantly differentially expressed in tumours showing low expression of GATA4 alone (G4^Lo^/G6^Hi^) (q-value<0.05). In contrast, in tumours showing only low GATA6 (G4^Hi^/G6^Lo^), 3914 genes (1871 up-and 2043 down-) were significantly dysregulated. Low expression of both genes (G4^Lo^/G6^Lo^) was associated with the most dramatic phenotype, with 6096 differentially expressed genes (2933 up-and 3163 down-) (Fig 2A).

**Figure 2.**
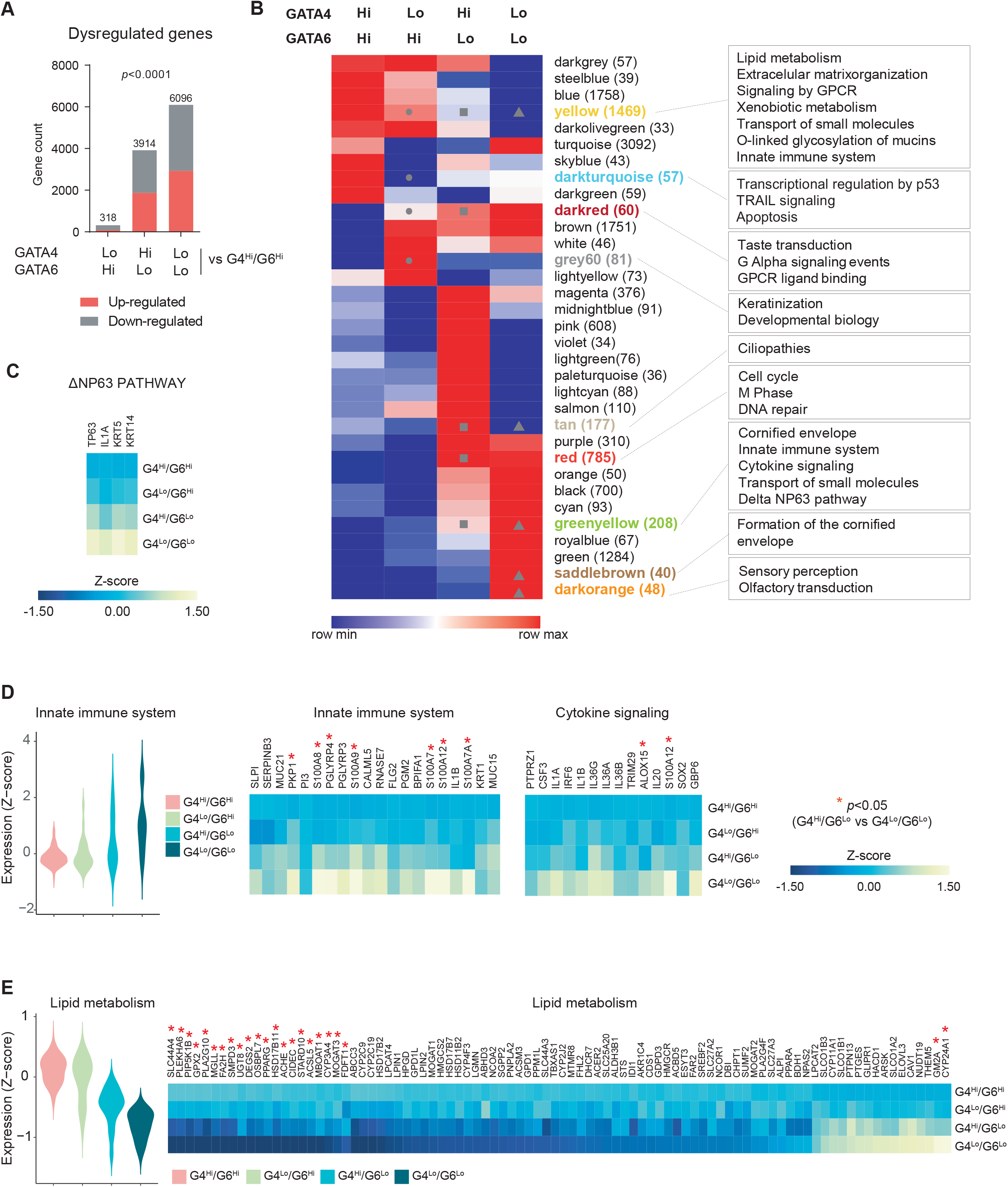
GATA4 down-regulation is associated with enhancement of the transcriptomic effect of GATA6 down-regulation in a module-dependent fashion. (A) Stacked bar plot representing the number of differentially regulated genes (q-value<0.05) (Chi-square test, *P<*0.0001). (B) Heatmap representing mean expression normalized by Z-score of genes contained in each module. Each gene programme is designated by color, numbers of genes per gene programme are shown in parentheses. Grey dot, square or triangle denote *P<*0.05 in G4^Lo^/G6^Hi^ vs G4^Hi^/G6^Hi^; G4^Hi^/G6^Lo^ vs G4^Hi^/G6^Hi^ and G4^Hi^/G6^Lo^ vs G4^Lo^/G6^Lo^ comparisons respectively (two-sided Mann–Whitney test). (C) Heatmap indicating expression of genes that overlap in the ΔNP63 pathway (PID) and the greenyellow module. (D) Violin plot and heatmap indicating expression of genes that overlap in Innate Immune System and Cytokine Signaling (REACTOME) signatures and the greenyellow module. (E) Violin plot and heatmap indicating expression of genes that overlap in the Lipid Metabolism (REACTOME) signature and the yellow module. Asterisk indicates *P<*0.05 when comparing G4^Hi^/G6^Lo^ vs G4^Lo^/G6^Lo^ samples (two-sided Mann–Whitney test).

To acquire insight into the direct or indirect effects of down-regulation of GATA4 and GATA6 we performed weighted gene correlation network analysis (WGCNA). This method identifies discrete groups of co-expressed genes and classifies them in modules (i.e. genetic programmes) by using the TOM algorithm[34]. These modules are often enriched for genes that share biological functions[35]. All tumours (n=247) clustered based on their Euclidean distance (Supplementary Figure 2A), excluding the presence of outliers. G4^Hi^/G6^Hi^ and G4^Lo^/G6^Hi^ samples distributed evenly across the dendrogram, whereas G4^Hi^/G6^Lo^ and G4^Lo^/G6^Lo^ tumours clustered together, in accordance with the results of the differential expression analysis.

To build a weighted gene network it is critical to choose a proper soft thresholding power (β) value. Thus, we screened for different soft thresholding powers and selected β = 4 for later analysis (Supplementary Figure 2B). Next, gene co-expression networks were generated corresponding to 33 co-expression modules (Supplementary Figure 2C,D). Modules were named after colors following WGCNA conventions[34].

#### GATA4-dependent programmes

We examined modules that were significantly dysregulated between G4^Lo^/G6^Hi^ and G4^Hi^/G6^Hi^ tumours by comparing the mean expression (normalized by Z-score) of the genes in each module (Figure 2B). Functional annotation of the grey60 module, selectively up-regulated in the G4^Lo^/G6^Hi^ group, revealed enrichment of genes involved in keratinization and development. The darkturquoise module, down-regulated in G4^Lo^/G6^Hi^ samples, contained genes related to apoptosis and/or regulated by p53.

#### GATA6-dependent programmes

We examined modules that were significantly dysregulated when comparing G4^Hi^/G6^Lo^ vs G4^Hi^/G6^Hi^ (describing the effect of low GATA6 expression alone). Only one gene programme (red) was identified, composed of 785 genes accounting for cell cycle, mitosis and DNA repair processes. Other GATA6-dependent modules following the same trend that did not reach significance are the magenta module, with genes involved in WNT signaling (e.g. *WNT7A, WNT7B, WNT10A, WNT11, FZD7, FZD2*), and the lightgreen module enriched in “Reactome_Interferon_Signaling”.

#### GATA4-dependent programmes in tumours with low GATA6 expression

We examined modules that were significantly dysregulated when comparing the G4^Hi^/G6^Lo^ group (GATA6 down-only) with the G4^Lo^/G6^Lo^ group. Functional annotation of gene programmes consistently up-regulated in the G4^Lo^/G6^Lo^ group revealed enrichment in processes related to formation of the cornified envelope and sensory perception in the saddlebrown and darkorange modules, respectively (Figure 2B, Supplementary Tables 3 and 4). Enrichment in processes related to the basal phenotype (i.e. “Formation of the cornified envelope”) was also found in the greenyellow module. Importantly, genes related to the ΔNP63 pathway were also contained in this module and were consistently up-regulated in the G4^Lo^/G6^Lo^ group (Figure 2C). In addition, immune pathways such as “cytokine signaling” and “innate immune pathways” (e.g. *IL1A, IL1B, IL36A*) were enriched in the greenyellow module, also up-regulated in the G4^Lo^/G6^Lo^ group (Figure 2D). The yellow module was significantly down-regulated in G4^Lo^/G6^Lo^ samples and was defined by genes involved in lipid metabolism, matrisome/extracellular matrix-related pathways, xenobiotic metabolism, O-linked mucin glycosylation, and inflammation. Reactome_“Metabolism_of_lipids” was the first annotation on the list with 91 overlapping genes (FDR q-value: 3.22E-21): 75/91 genes were down-regulated and 16/91 were up-regulated in the G4^Lo^/G6^Lo^ group (Figure 2E). Among the former are those related to fatty acid metabolism (e.g. *FA2H, MGLL*), cholesterol biosynthesis (e.g. *FDFT1*), and acetyl-choline metabolism (e.g. *ACHE, ACSL5*). *GM2A* and *CYP24A1* are the most significant lipid-related genes up-regulated in G4^Lo^/G6^Lo^ samples.

### Concomitant low expression of GATA6 and GATA4 is associated with the worst patient outcome

To interrogate whether the molecular cooperation described above has a clinical translation, we evaluated GATA4 and GATA6 protein expression in a multicenter TMA study of PDAC samples from patients undergoing tumour resection (n=745). This TMA series includes both patients enrolled in a clinical trial of adjuvant chemotherapy (ESPAC-3 cohort)[32] and a more diverse group of patients receiving standard therapy[29–31]. Table 1 summarizes the characteristics of the patients included in the study. We first compared the clinical features of the various patient series, focusing on outcome. Median overall survival was highest for patients from the ESPAC-3 study (23.5 months vs 14.5, 17.6, and 12.4 for the other series) (Supplementary Figure 3A). Considering all patients together, median survival was higher for patients who had received adjuvant chemotherapy: 5-FU (23.5 months), gemcitabine (22.8 months), or unknown (22.0 months) (differences not being significant) vs those who had not (14.6 months) (Supplementary Figure 3B). Similar survival was observed among patients from the ESPAC-3 trial and those receiving adjuvant therapy in the conventional setting (not shown).

**Table 1.**
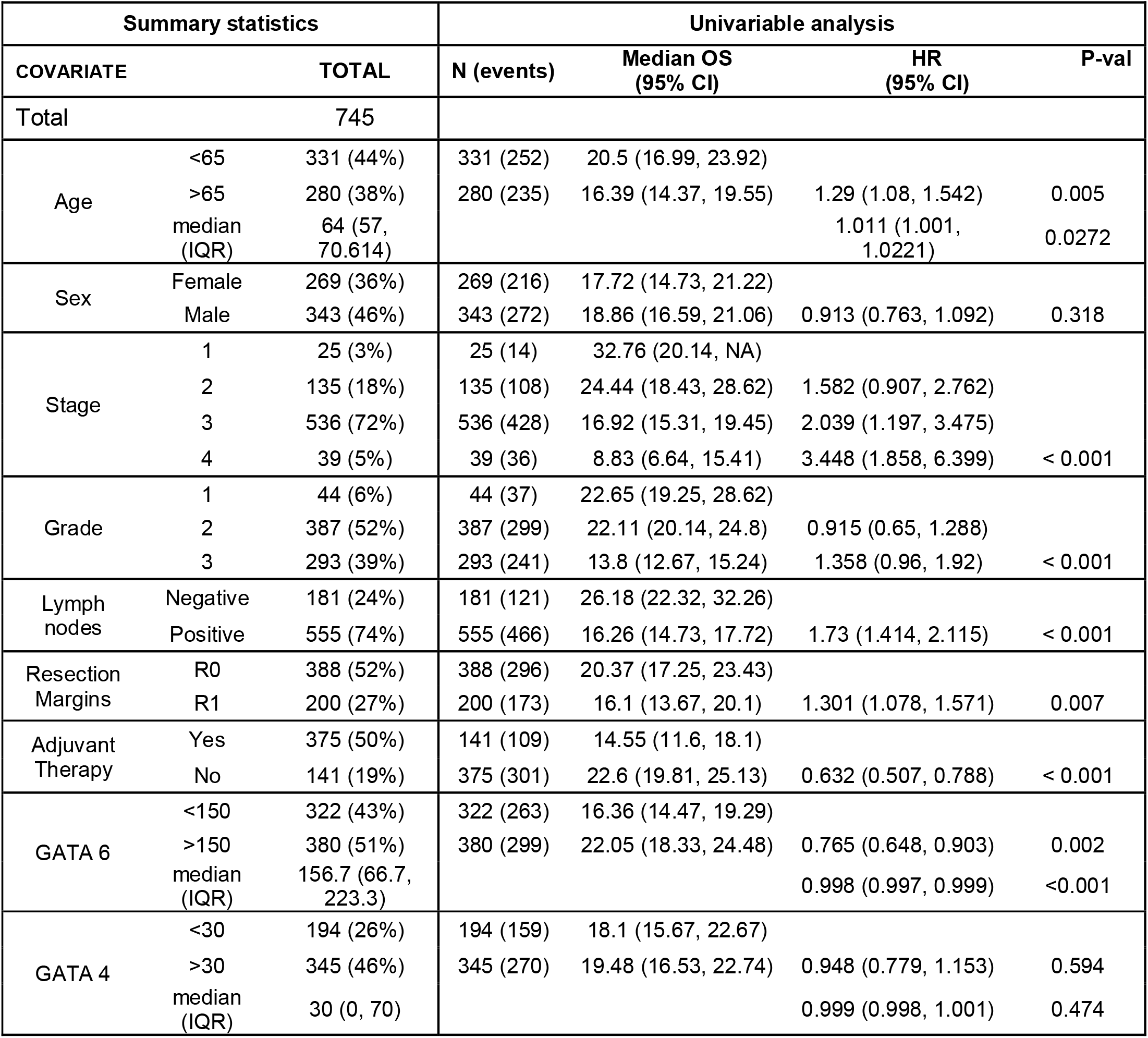
Baseline demographics, clinical-pathological characteristics and expression of GATA4 and GATA6 in tumours from patients included in the study. Results of the univariable analysis of survival.

| Summary statistics |  |  | Univariable analysis |  |  |  |
| --- | --- | --- | --- | --- | --- | --- |
| COVARIATE |  | TOTAL | N (events) | Median OS (95% CI) | HR (95% CI) | P-val |
| Total |  | 745 |  |  |  |  |
| Age | <65 | 331 (44%) | 331 (252) | 20.5 (16.99, 23.92) |  |  |
|  | >65 | 280 (38%) | 280 (235) | 16.39 (14.37, 19.55) | 1.29 (1.08, 1.542) | 0.005 |
|  | median (IQR) | 64 (57, 70.614) |  |  | 1.011 (1.001, 1.0221) | 0.0272 |
| Sex | Female | 269 (36%) | 269 (216) | 17.72 (14.73, 21.22) |  |  |
|  | Male | 343 (46%) | 343 (272) | 18.86 (16.59, 21.06) | 0.913 (0.763, 1.092) | 0.318 |
| Stage | 1 | 25 (3%) | 25 (14) | 32.76 (20.14, NA) |  |  |
|  | 2 | 135 (18%) | 135 (108) | 24.44 (18.43, 28.62) | 1.582 (0.907, 2.762) |  |
|  | 3 | 536 (72%) | 536 (428) | 16.92 (15.31, 19.45) | 2.039 (1.197, 3.475) |  |
|  | 4 | 39 (5%) | 39 (36) | 8.83 (6.64, 15.41) | 3.448 (1.858, 6.399) | < 0.001 |
| Grade | 1 | 44 (6%) | 44 (37) | 22.65 (19.25, 28.62) |  |  |
|  | 2 | 387 (52%) | 387 (299) | 22.11 (20.14, 24.8) | 0.915 (0.65, 1.288) |  |
|  | 3 | 293 (39%) | 293 (241) | 13.8 (12.67, 15.24) | 1.358 (0.96, 1.92) | < 0.001 |
| Lymph nodes | Negative | 181 (24%) | 181 (121) | 26.18 (22.32, 32.26) |  |  |
|  | Positive | 555 (74%) | 555 (466) | 16.26 (14.73, 17.72) | 1.73 (1.414, 2.115) | < 0.001 |
| Resection Margins | R0 | 388 (52%) | 388 (296) | 20.37 (17.25, 23.43) |  |  |
|  | R1 | 200 (27%) | 200 (173) | 16.1 (13.67, 20.1) | 1.301 (1.078, 1.571) | 0.007 |
| Adjuvant Therapy | Yes | 375 (50%) | 141 (109) | 14.55 (11.6, 18.1) |  |  |
|  | No | 141 (19%) | 375 (301) | 22.6 (19.81, 25.13) | 0.632 (0.507, 0.788) | < 0.001 |
| GATA 6 | <150 | 322 (43%) | 322 (263) | 16.36 (14.47, 19.29) |  |  |
|  | >150 | 380 (51%) | 380 (299) | 22.05 (18.33, 24.48) | 0.765 (0.648, 0.903) | 0.002 |
|  | median (IQR) | 156.7 (66.7, 223.3) |  |  | 0.998 (0.997, 0.999) | <0.001 |
| GATA 4 | <30 | 194 (26%) | 194 (159) | 18.1 (15.67, 22.67) |  |  |
|  | >30 | 345 (46%) | 345 (270) | 19.48 (16.53, 22.74) | 0.948 (0.779, 1.153) | 0.594 |
|  | median (IQR) | 30 (0, 70) |  |  | 0.999 (0.998, 1.001) | 0.474 |

GATA4 and GATA6 protein expression levels were also dichotomized as described for mRNA expression. GATA6 histoscores were distributed more evenly across samples than GATA4 histoscores, the distribution of which was skewed towards lower values (Figure 3A). A moderate, significant, correlation between the expression of both proteins was observed (rho=0.31, p-value = <2.2e-16; Spearman’s test) (Figure 3B). Similar results were obtained when testing each individual cohort separately (Supplementary Figure 4A-H). Representative immunohistochemical results are shown in Figure 3C.

**Figure 3.**
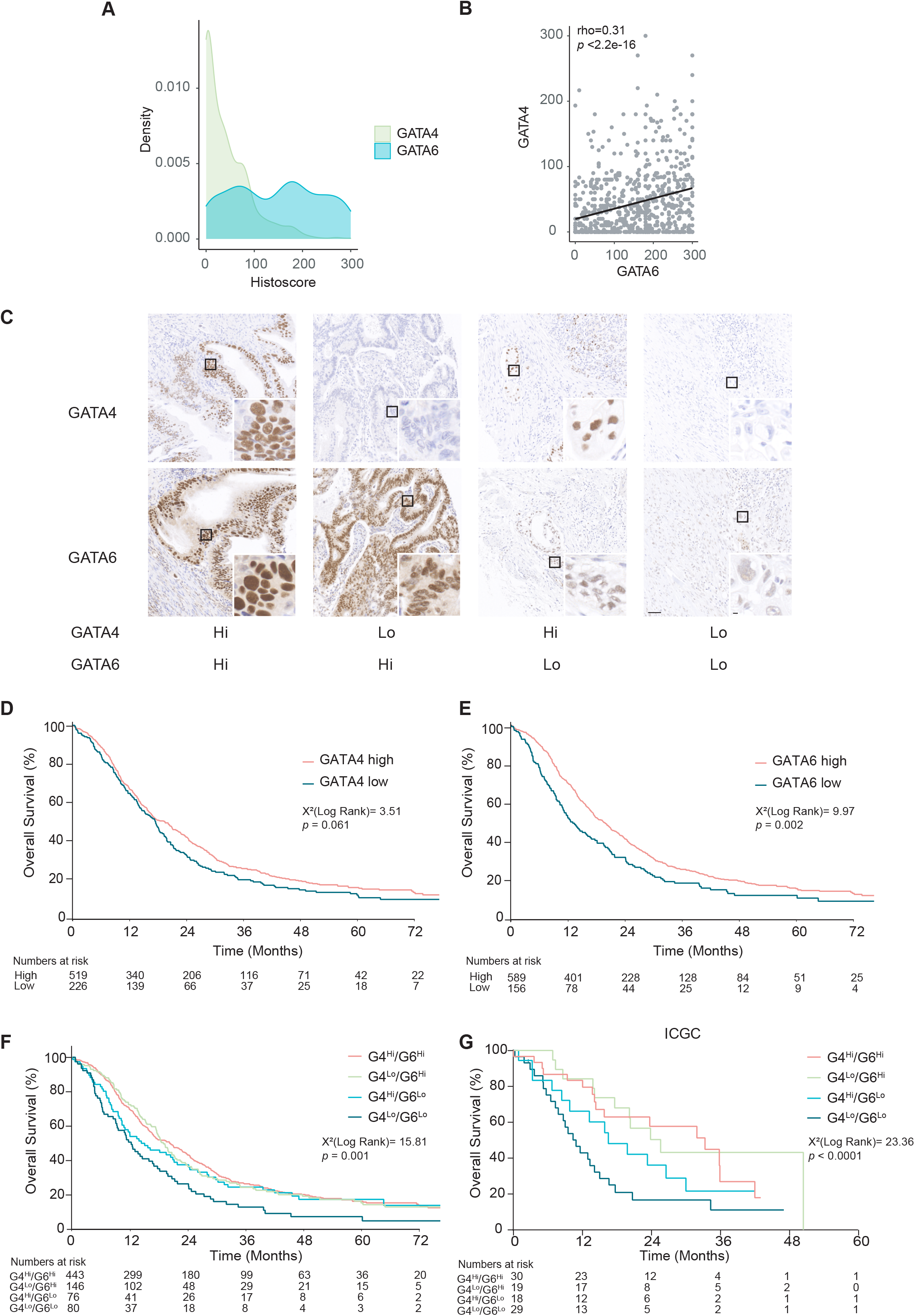
Patients whose tumours express low levels of both GATA4 and GATA6 have the worse outcome. (A) Density plot showing GATA4 and GATA6 histoscore distribution in all cases (n=745). (B) Scatter plot showing a positive correlation between GATA4 and GATA6 histoscore across all samples (Spearman correlation). (C) Representative images showing GATA4 and GATA6 expression, detected by IHC, in tumour samples from each of the combined expression categories. Scale bar: 50uM. Magnification, scale bar: 5uM. (D) Kaplan-Meier plot of overall survival by GATA4 IHC status (n=745). (E) Kaplan-Meier plot of overall survival by GATA6 IHC status (n=745). (F) Kaplan-Meier plot showing the association of combined GATA4 and GATA6 IHC status with survival (n=745). (G) Kaplan-Meier plot comparing survival of patients from ICGC-Bailey et al.[11] according to the expression of GATA4 and GATA6 mRNA (n=96).

Survival analyses showed that GATA4 expression, as a dichotomous variable, was not associated with overall survival [hi vs low, 19.4 (95% CI 16.6-22.1) vs 17.5 (14.6-19.3)] (Figure 3D). In contrast, GATA6 expression was significantly associated with a shorter overall survival [hi vs low, 19.7 (17.4-21.6) vs 13.0 (10.9-17.6), p value<0.001] (Figure 3E). Patients with tumours showing low expression of both GATA4 and GATA6 had the worst outcome (G4^Lo^/G6^Lo^ vs all others, 12.2 (9.3-19.9) vs 19.3 (17.3-21.0), p value<0.001) (Figure 3F). When the results of each series were analyzed separately, the same trend was observed but the differences were not statistically significant, likely due to the smaller sample size (Supplementary Figure 5A-D). Similar results were obtained when the same strategy was applied to the transcriptomic data from the Australian Pancreatic Cancer Initiative (Figure 3G).

Univariable analysis of all factors included in the dataset was performed (Table 1). Age, stage, grade, lymph node involvement, positive resection margins, adjuvant therapy, and GATA6 as a single marker were all significantly associated with outcome. For the multivariable analysis, all the clinical/pathological terms were considered in the final model, selected using a backwards stepwise selection. In addition, the combined expression of GATA4/GATA6 was included. Grade, lymph node involvement, adjuvant therapy, resection margins, and low expression of both GATA proteins were significantly and independently associated with survival. The HR for low expression of both GATAs, when compared against any other configuration of protein expression, was 1.59 (95%CI 1.22-2.10; *p<*0.001) (Table 2). Exploratory multivariable analyses suggest that the impact of GATA4/6 is consistent across all levels of the adjuvant therapy covariate (not shown); the interaction term between GATA4/6 status and adjuvant therapy was non-significant (Supplementary Table 5 and Supplementary Figure 6). These findings reveal that combined GATA4/GATA6 expression identifies the group of resectable PDAC patients with the worst outcome.

**Table 2.** Multivariable modeling of overall survival in the patients included in the study.

| Covariate | Level | Estimate (se) | HR (95% CI) | P-value |
| --- | --- | --- | --- | --- |
| Grade | Poor |  |  |  |
|  | Moderate | -0.31 (0.093) | 0.73 (0.612, 0.88) | 0.001 |
|  | Well | -0.21 (0.181) | 0.81 (0.569, 1.157) | 0.248 |
| Stage | 1 |  |  |  |
|  | 2 | 0.26 (0.293) | 1.3 (0.733, 2.313) | 0.368 |
|  | 3 | 0.33 (0.286) | 1.39 (0.791, 2.427) | 0.255 |
|  | 4 | 0.85 (0.337) | 2.33 (1.206, 4.514) | 0.012 |
| Nodes | Negative |  |  |  |
|  | Positive | 0.49 (0.111) | 1.63 (1.309, 2.022) | <0.001 |
| Adjuvant therapy | Yes |  |  |  |
|  | No | 0.46 (0.202) | 1.58 (1.064, 2.348) | 0.023 |
|  | Unknown | 0.51 (0.202) | 1.66 (1.12, 2.469) | 0.012 |
| Resection margins | No |  |  |  |
|  | Yes | 0.29 (0.104) | 1.34 (1.091, 1.638) | 0.005 |
|  | Unknown | -0.11 (0.261) | 0.9 (0.538, 1.499) | 0.681 |
| GATA 4/6 | Other |  |  |  |
|  | Lo-Lo | 0.47 (0.14) | 1.59 (1.22, 2.10) | 0.001 |

### Low GATA4 and GATA6 expression is associated with increased liver metastasis

To assess GATA4 and GATA6 expression during tumour progression, we first assessed protein expression of GATA4 and GATA6 -and a panel of well-established classical and basal markers -in a small cohort of 6 matched pairs of primary PDAC and distant metastasis (see methods), as well as 2 additional metastases. We classified samples in four groups according to GATA4 and GATA6 median histoscore. All primary tumours tested along with their paired metastasis had a high expression of CDH-1 and most of them lacked the expression of the tested basal markers (Figure 4A).

**Figure 4.**
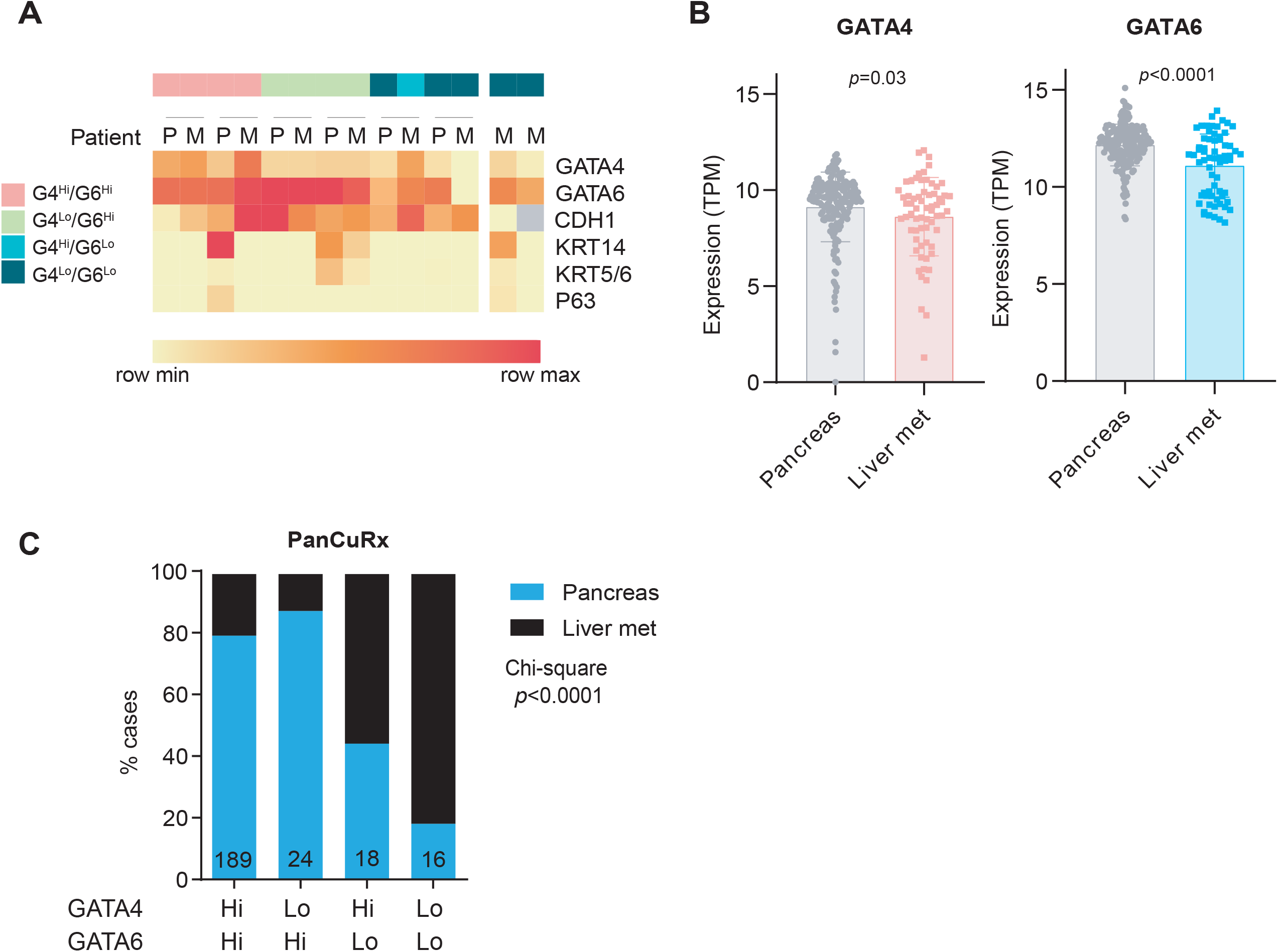
Down-regulation of GATA4 and GATA6 is associated with increased liver metastasis. (A) Heatmap representing relative histoscore of the indicated genes per paired sample. P= primary tumour, M=metastatic sample. (B) GATA4 and GATA6 expression in primary and metastatic samples from the PanCuRx dataset (Two-sided Mann–Whitney test). (C) Stacked bar plot representing the distribution of primary tumours and metastatic samples according to GATA4/GATA6 expression categories (Chi-square test, *P<*0.0001).

Leveraging on the PanCuRx dataset, which includes a large number of metastatic samples, we compared expression of GATA4, GATA6, and the basal and classical markers in primary and metastatic tumours. We observed a significant positive correlation between GATA6 and CDH1 in metastatic samples while no significant correlation was observed with GATA4 (Supplementary Figure 7A). In the case of the basal markers KRT5/6, KRT14 and TP63, we observed a consistent significant negative correlation with both GATA factors in metastatic samples, also less significant in primary tumours (Supplementary Figure 7B-D). These findings confirm that GATA4 and GATA6 expression is inversely associated with the basal phenotype in metastatic samples.

We then compared GATA4 and GATA6 levels in primary vs metastasis and, as expected, we observed a significantly lower expression of both genes in the metastases (Figure 4B). Interestingly, 50% of samples in the G4^Hi^/G6^Lo^ group and 80% of the G4^Lo^/G6^Lo^ samples were liver metastases (Figure 4C).

## DISCUSSION

In PDAC, GATA6 is a marker and a master regulator of the classical subtype. Here, we provide strong evidence that GATA4 plays an important role in the absence of GATA6. The effect of GATA4 on PDAC biology is reflected on patient outcome.

GATA transcription factors bind with high affinity to consensus GATA sites and have very similar binding specificities. A challenge in the understanding of their biology is the dissection of their overlapping and unique functions. In some cases, selective expression of one vs other GATA genes accounts for tissue-specific functions. This is the case of GATA6 in peritoneal macrophages[36] or of GATA4 in senescence in fibroblasts[37]. When multiple GATA proteins are expressed in a given cell, dissecting gene-specific functions is more challenging. Genetic mouse models have shown that deletion of *Gata4* or *Gata6* does not impact on pancreatic development, while deletion of both genes blunts pancreatic proliferation and differentiation[23,24]. A study using human pluripotent stem cells has reported the relevance of GATA4 dosage in a GATA6 heterozygous context, highlighting the interactions between both factors[38]. In the intestine, inactivation of *Gata4* and *Gata6* unveils a time-and space-constrained redundancy[39]. The complexity underlying their cooperation has been revealed in murine colorectal cancer, where *Gata6* deletion in adenomas results in increased BMP levels and a blockade of tumour stem cell self-renewal[40]. Genetically-engineered expression of one of the proteins under the regulatory elements of the paralogue gene has shown that GATA4 is required for liver and heart development[41].

The cooperativity between GATA proteins in cancer has been poorly explored. In PDAC, GATA6 has been widely studied but there are few, inconclusive, reports on GATA4 expression and activity[42–45]. *GATA6* is amplified and overexpressed in a subset of pancreatic and hepatobiliary tumours[11,12,46,47]. In contrast, *GATA4* copy number changes are infrequent. We hypothesized that the combined analysis of GATA4 and GATA6 expression might reveal specific biological functions and relevant clinical outcome associations. Of note, GATA6 mRNA levels have been shown to accurately correlate with protein levels in human PDAC[16]. RNA-Seq analysis of tumours from the PanCuRx series using an agnostic module identification strategy revealed that GATA6 down-regulation has the major transcriptomic impact; tumours displaying only low GATA4 levels had a molecular profile similar to those with preserved expression of both transcription factors. Dependency on GATA4 and GATA6 has been reported in gastric cancer *in vitro* upon knockdown of one or both genes, either directly or indirectly through other transcription factors, including CDX2[48].

Modules with higher activity in tumours with combined down-regulation of GATA4 and GATA6 were enriched in “basal-ness”-related genes and in the ΔNp63 pathway. GATA6 down-regulation is necessary, but not sufficient, to activate the expression of the ΔNp63 programme and the basal phenotype both in mouse and patient tumours. The cooperative effect resulting from the absence of both GATA proteins is similar to that observed when GATA6 is down-regulated concomitantly with HNF1A or HNF4[49]. These pathways likely contribute to the aggressiveness of the Basal tumours by favouring an epithelial-to-epithelial transition that precedes an EMT. Genes in the cytokine signaling and immune pathways were also enriched in G4^Lo^/G6^Lo^ tumours, possibly contributing to immune escape and metastasis[49]. GATA4 and GATA6 levels were lower in liver metastases and they negatively correlate with basal markers. Comparison of G4^Hi^/G6^Lo^ vs G4^Lo^/G6^Lo^ metastatic samples identified CDX2 among the top 5 down-regulated genes in G4^Lo^/G6^Lo^ samples (not shown).

G4^Lo^/G6^Lo^ tumours also showed a down-regulation in genes involved in cholesterol and fatty acid biosynthesis. Metabolic profiling and bioinformatic analyses have revealed two main PDAC metabolic subtypes: cholesterogenic and glycolytic. The former is associated with the classical subtype and with a better prognosis, while the latter is associated with the basal subtype[50,51]. Accordingly, using previously described signatures[51], we found a significant up-regulation of the glycolysis pathway and down-regulation of the cholesterol biosynthesis pathway in the G4^Lo^/G6^Lo^ tumours (not shown). Disruption of cholesterol synthesis favors a switch to a basal phenotype that is mediated by up-regulation of TGF-beta[52]. Several membrane small metabolite transporters are expressed at higher levels in classical than in basal tumours; among them is the cholesterol transporter NPC1L1, the pharmacological and genetic inhibition of which can suppress the growth of PDAC cells in vitro and in mice[53]. The transcriptional cooperation resulting from the down-regulation of both GATA proteins supports the acquisition of aggressive tumour features.

Our study was prompted by the clinical value of GATA6 as a marker of outcome and therapeutic stratification. In the largest patient series reported until now, we show that patients with tumours showing low levels of GATA6, but not of GATA4, have poor outcome but patients with tumours expressing low levels of both proteins had the worse survival. The association with outcome was significant as an independent predictor in the multivariable analysis and the magnitude of the effects -while modest -is similar to that of the other important outcome predictors such as lymph node involvement, administration of adjuvant therapy, and positive resection margins. The association was consistent across patient cohorts involving both patients selected for trials and those treated in the standard clinical setting. The large sample size of our study supports the robustness of the observations.

Future studies should consider the use of full sections instead of TMA cores to account for cellular heterogeneity. *GATA6* and *GATA4* are among the genes with higher 5hmC density in circulating cell free DNA in patients with PDAC[54]. 5hmC is associated with transcription and its cfDNA profiles mirror the hydroxy-methylation profile in tumour samples, suggesting that it may be possible to subtype patients using non-invasive methods to select optimal therapy.

The robustness of the IHC assays, already confirmed by independent investigators in the case of GATA6[55], calls for validation of our findings in prospective studies and the extension to patients with advanced PDAC. It will also be important to determine whether the combined assessment of GATA6 and GATA4 improves prediction of response to chemotherapy.

## Acknowledgements

We thank Peter Bailey, Eduard Batlle, Adrià Caballe, Mark Kalisz, Bernhard Kloesch, Miriam Marqués, Jaime MartÍnez de Villarreal, Faiyaz Notta, and Camille Stephan-Otto Atolini for valuable contributions and the Histopathology Unit of CNIO for technical help.

## Funding

The work was supported, in part, by the following grants: RTI2018-101071-B-I00 (Ministerio de Ciencia, Innovación y Universidades-MCIU, Madrid, Spain) to FXR, PI18/01347 (ISCIII-FIS, Madrid, Spain) to NM, PRECODE (EU-MSCA: 861196) to CP. MPdA was supported by a Ph.D. Fellowship from MCIU. CNIO is supported by MCIU as a Centro de Excelencia Severo Ochoa (grant SEV-2015-0510).

## Competing interests

The authors declare no conflict of interest.

## Author contributions

MPdA and FXR, together with PM, designed the study, analyzed the results, and wrote the manuscript. MPdA performed the experiments and bioinformatics analyses. RJ performed the statistical analysis with contributions from NM. CP, AMS, TK, PR, and WW provided samples and clinical information. EC, WG, PG, DP, MB, TH, and JPN were involved in the ESPAC-3 trial and provided samples and clinical information. All authors provided comments to the manuscript. FXR supervised the overall conduct of the study and obtained funds.

## Notes

### Competing Interest Statement

The authors have declared no competing interest.

