## Supplementary Materials for "GATA4 and GATA6 loss-of-expression is associated with extinction of the classical programme and poor outcome in pancreatic ductal adenocarcinoma"

**M. P. de Andrés et al.**

### SUPPLEMENTARY MATERIAL

#### Supplementary Methods and Materials

**Expression datasets.** Quantile-normalized read counts of 228 samples of normal pancreas from the GTEx version 8 dataset (GTEx_Analysis_2017-06-05_v8_RNASeQCv1.1.9_gene_tpm.gct.gz) were retrieved from https://gtexportal.org/home/datasets. The following PDAC datasets were downloaded from the indicated source and used in the analyses: ICGC (Bailey et al) [RNA-seq, bulk tissue (qPure score 31-99), Log2 (count per million) normalization, n=96, supplementary table from publication[11]], PanCuRx (RNA-seq, microdissected tissue, TPM normalization, n=247, EGAS00001002543).

**Tumour classification**. To establish tumour categories, the distribution of GATA4 and GATA6 expression values was assessed. In the PanCuRx dataset, a bimodal distribution was observed; samples with GATA4 expression <7.6 transcripts per million (TPM) (16%) were considered GATA4 low and the rest was considered GATA4 high. Samples with GATA6 expression <10.3 TPM (14%) were considered GATA6 low and the rest was considered GATA6 high. In the Bailey et al. dataset, a normal distribution was observed. Patients were stratified according to the median value of GATA4 or GATA6 expression. Samples above the median value were considered (GATA4 or GATA6) high and, conversely, samples under the median value were considered (GATA4 or GATA6) low. Based on each of these stratification approaches, four categories were defined in both datasets: GATA4 high/GATA6 high ($G4^{Hi}/G6^{Hi}$), GATA4 low/GATA6 high ($G4^{Lo}/G6^{Hi}$), GATA4 high/GATA6 low ($G4^{Hi}/G6^{Lo}$), and GATA4 low/GATA6 low ($G4^{Lo}/G6^{Lo}$).

**Differential gene expression analysis.** Protein-coding genes were filtered from the PanCuRx dataset, including 19,618 genes in the analysis. Differential gene expression analysis was performed using Comparative Marker Selection (version 10.1) and Comparative Marker Selection Viewer (version 9) tools[26] through GenePattern using the default parameters.

**Weighted gene correlation network analysis (WGCNA)**. The R package WGCNA was applied to analyze the PanCuRx dataset. Only protein-coding genes were selected (n=19,618). First, 151 genes were excluded from the calculation using "goodSamplesGenes" function. Next, the appropriate soft-thresholding power (β), used to detect co-expression similarity and calculate adjacency, was screened and selected by using the ''pickSoftThreshold'' function (β = 4). For network construction, Pearson's correlation coefficients were calculated based on co-expression similarity for each pair of genes. The resulting Pearson's correlation matrix was transformed into an adjacency matrix using a power function (β = 4). The topological overlap matrix (TOM) algorithm was used to identify modules of densely interconnected genes [27]. The minimum number of genes per module was set as 30. To strengthen the reliability of the modules, modules bearing <0.45 were merged. The grey module contains genes that were not assigned to specific modules.

**Gene Set Enrichment Analysis (GSEA).** Identification of biological processes in gene modules was performed using the tool "Investigate Gene Sets" provided by the Molecular Signatures Database v7.4 [28]. Gene sets contained in Collection C2 were used (BIOCARTA, KEGG, PID, REACTOME and WikiPathways). Only gene sets with FDR q-value <0.05 were considered.

**Patient cohorts and clinical information.** The study design included the analysis of multiple patient cohorts from the outset in order to increase the statistical precision of the analyses and the representativity of the findings. All study sites provided tissue microarrays (TMAs) containing tumour samples from patients who underwent surgical resection. Informed consent was obtained from all patients participating in the study. A summary of relevant information is provided in Supplementary Table 1.

*Cohort 1 (Berlin).* Tissue microarrays were assembled from pancreatic resection specimen obtained between 1991 and 2006 at the Charité University Hospital (Berlin, Germany). The use of this tumour cohort for biomarker analysis has been approved by the Institutional Review Board (IRB; ethics committee) of the Charité University (EA1/06/2004) [29].

*Cohort 2 (Munich).* Tissue microarrays were assembled from pancreatic resection specimens obtained between 2007 and 2011 at the Department of Surgery, Klinikum rechts der Isar, TU München, Germany[30]. The study was approved by the ethics committee of the TU München, Germany (documents no. 1926/2007 and 126/2016S).

*Cohort 3 (Regensburg).* Tissue microarrays were prepared from patient samples obtained after appropriate informed consent in Dresden (Institute of Pathology, University Hospital Dresden), Regensburg (Institute of Pathology, University Hospital Regensburg) and Jena (Institute of Pathology, University Hospital Jena). Samples were collected from 1993 to 2009 [31]. The study was approved by the Ethikkommission an der Technischen Universität Dresden (ref. 170_16 B).

*Cohort 4 (ESPAC-3).* These samples were part of the ESPAC-3 trial, a two arm, international, open-label, phase 3, randomized controlled trial comparing adjuvant fluorouracil plus folinic acid versus gemcitabine following curative resection of PDAC [32]. The study was approved by the national committees of all the participating countries as outlined in the original publication. ESPAC-T project research is covered under IRAS project ID 231188) from the North Western-Liverpool Central Research Ethics Committee. The results of GATA6 expression in this dataset have been previously reported [3].

*Paired Primary-Metastasis cohort.* Tissue microarrays of a small cohort of primary PDACs and matched distant metastases from pancreatic resection specimen and distant metastases obtained between 2004 and 2020 at the Klinikum rechts der Isar, TU München were assembled. The use of the tumor cohort for biomarker analysis has been approved by the ethics committee of the Technical University München, Germany.

**Immunohistochemical analysis.** Sections of formalin-fixed paraffin-embedded tissue microarray blocks were heated for 30 min at 55 ºC, deparaffinized with xylol, and rehydrated with an alcohol series and distilled water. Sections were boiled in 10mM citrate buffer (pH=6.0) for 10 min for antigen retrieval. Next, sections were incubated for 30 min with 3% $H_2O_2$ in methanol, then washed with PBS, and incubated for 1h with 2%

BSA in PBS at room temperature. Primary antibodies were added overnight at 4 ºC in 2% BSA in PBS. The following primary antibodies were used: GATA6 (R&D systems, AF1700, 0.2ug/mL), GATA4 (R&D systems, MAB2606, 1ug/mL). Slides were washed with PBS/0.1% Triton X-100 and, only for GATA6 staining, they were incubated with polyclonal rabbit anti-goat HRP (Dako, 1:200), for 1h at room temperature. Finally, slides were washed three times with PBS/0.1% Triton X-100 and incubated with the corresponding EnVision+ HRP secondary antibodies (Dako). DAB was used as chromogen and nuclei were counterstained for 1'30'' with Carazzi's Hematoxylin. P63, CDH1, KRT5/6 and KRT14 stainings were performed in an automated immunostaining platform (Discovery XT, Ventana, Roche), using the following primary antibodies: mouse monoclonal anti-P63 (4A4, ready to use, Roche, 790-4509), mouse monoclonal anti-CDH1 (36, 1/1000, Becton Dickinson, 610182), rabbit polyclonal anti-KRT14 (1/4500, Covance, PRB-155P) and rabbit polyclonal anti-KRT5/6 (1/5000, Covance, PRB-160P).

**Histoscore quantification.** To determine histoscore (HS), the proportion of reactive tumour cells was estimated and multiplied by the intensity of the staining (from 0-null to 3-strong). The obtained HS ranged from 0 to 300. If more than one tumour core was available from a given case, the mean value was used. In case of major discrepancies of histoscores, the slides were re-reviewed to confirm the appropriateness of the scorings. All samples were reviewed blindly by two researchers involved in the study (M.P.d.A. and F.X.R.).

**Survival analysis.** The primary outcome was overall survival (OS), measured as the time elapsed from surgery until death by any cause. Patients are censored at the date last known alive. Information was retrieved from 4 sources corresponding to the different TMA series. Data were retained for analysis only if expression information was available for both GATA4 and GATA6. Continuous data were summarized as median (IQR) and categorical data were summarized as frequencies of counts with associated percentages. Estimates of OS were obtained using Kaplan Meier curves. The prognostic value of GATA4 and GATA6 expression was explored graphically by dichotomizing into two-level high/low categorical covariates to provide comparisons based on the bottom 20% and top 80% expressing cases to differentiate between no/low activity vs moderate/high activity. Covariates were dichotomized within each data source to allow for heterogeneity. The prognostic impact of explanatory covariates was evaluated using multivariable Cox proportional hazards models with results presented as hazard ratios (HR) with 95% confidence intervals (CI). Clinical and pathological variables, as well as data source, were included for modelling. All covariates were considered for inclusion with model selection performed using a backwards step-wise procedure based on Akaikes Information Criterion. GATA4 and GATA6 were considered as both main effects and as two-level interactions. Models were explored including GATA4 and GATA6 as both continuous and categorical covariates to assess consistency of conclusions. A p-value $< 0.05$ was used to determine statistical significance throughout. All analyses were performed using R (Version 4).

#### SUPPLEMENTARY FIGURES

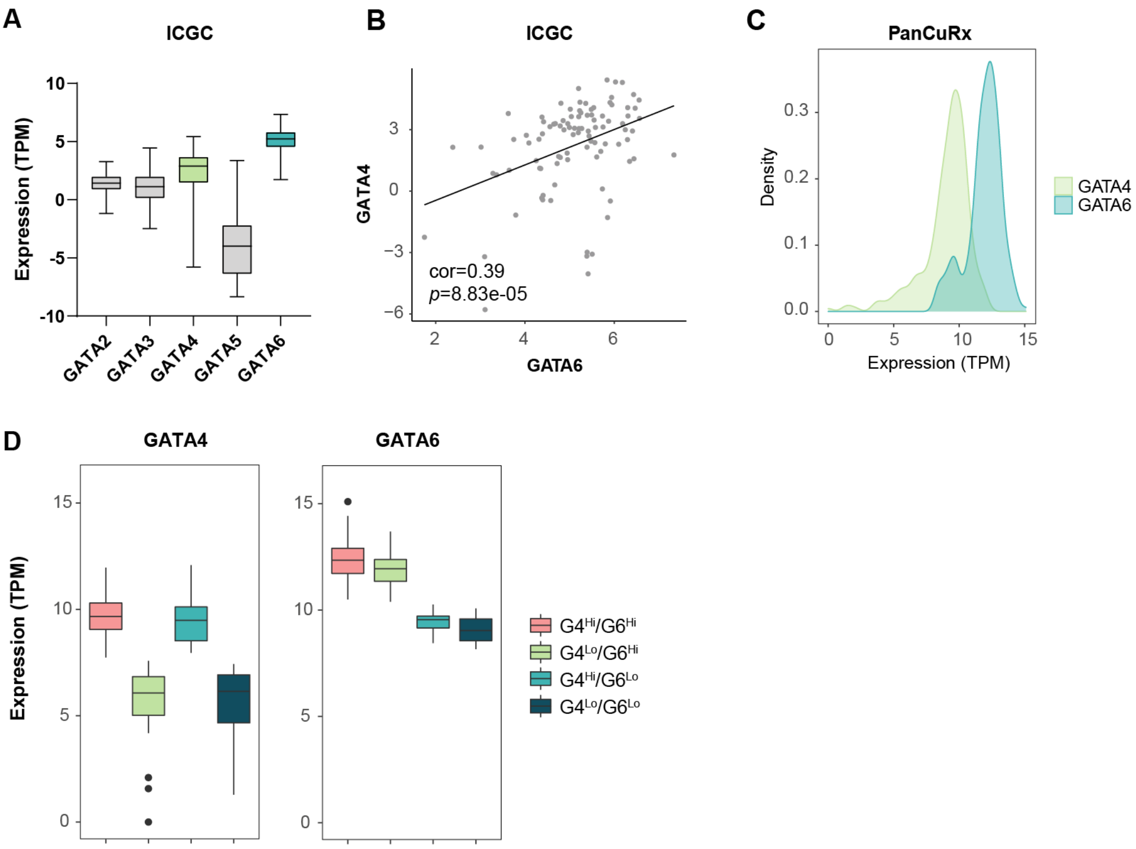


**Supplementary Figure 1. GATA4 and GATA6 expression in PDAC samples from the ICGC dataset.** (A) mRNA expression of GATA1-6 in the ICGC dataset of human PDAC (ref). Boxplot is annotated by a Kruskal-Wallis P value. (B) Scatter plot showing a positive correlation between GATA4 and GATA6 mRNA expression levels in the ICGC dataset (Pearson correlation). (C) Density plot showing the distribution of GATA4 and GATA6 mRNA expression levels in the PanCuRx dataset. (D) Boxplot showing GATA4 and GATA6 levels according to GATA4/GATA6 expression categories in the PanCuRx dataset.

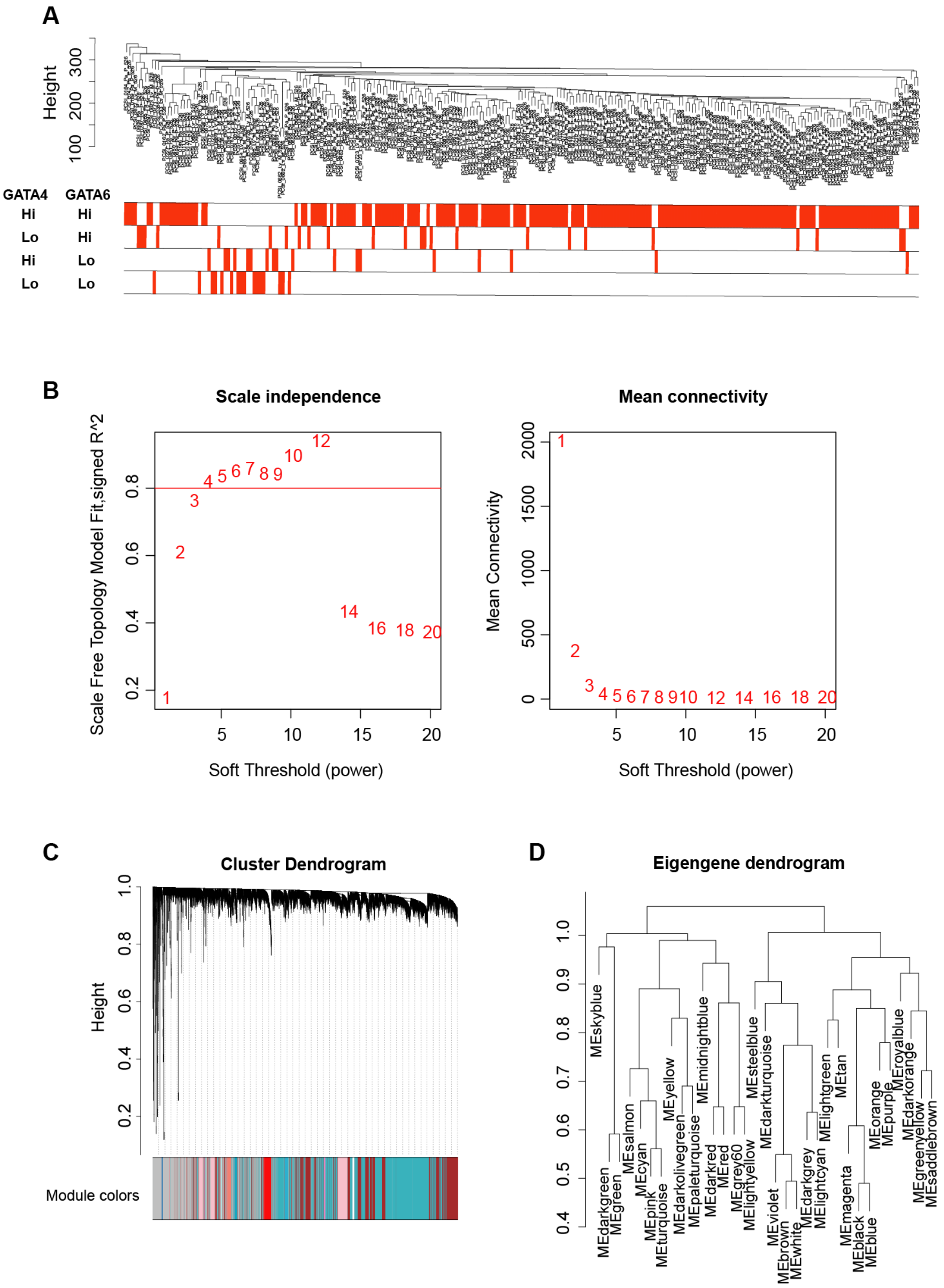


**Supplementary Figure 2. WGCNA identified 33 co-expression modules**. (A) Dendrogram showing sample clustering to detect outliers. Sample distribution across GATA4/GATA6 expression categories was assessed. Every red line corresponds to an individual sample. (B) Screening for soft-thresholding powers and analysis of the mean connectivity for various soft-thresholding powers. (C) Hierarchical clustering dendrogram of genes for identifying consensus modules. (D) Clustering dendrograms of consensus module eigengenes (gene programmes).

**A**

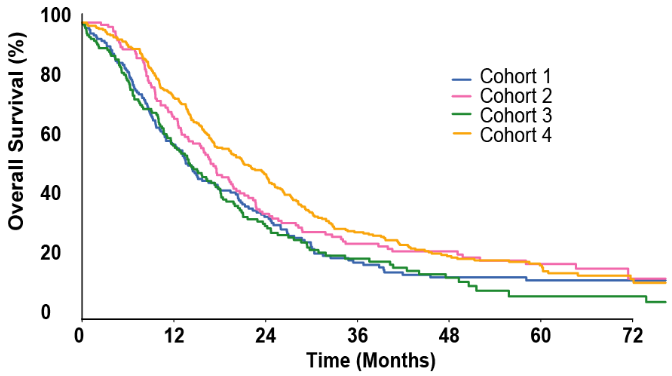


Numbers at risk

| | | | | | | | |
|---|---|---|---|---|---|---|---|
| Cohort 1 | 165 | 92 | 49 | 24 | 15 | 13 | 10 |
| Cohort 2 | 133 | 90 | 46 | 32 | 22 | 15 | 4 |
| Cohort 3 | 165 | 89 | 41 | 20 | 11 | 4 | 4 |
| Cohort 4 | 282 | 208 | 136 | 77 | 48 | 28 | 11 |

**B**

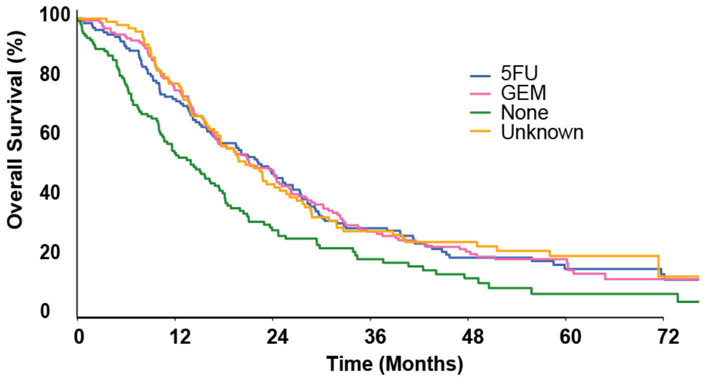


Numbers at risk

| | | | | | | | |
|---|---|---|---|---|---|---|---|
| 5FU | 132 | 94 | 62 | 37 | 21 | 12 | 8 |
| GEM | 150 | 114 | 74 | 40 | 27 | 16 | 3 |
| None | 141 | 71 | 31 | 16 | 9 | 3 | 3 |
| Unknown | 93 | 72 | 41 | 24 | 17 | 11 | 2 |

**Supplementary Figure 3. Survival analysis of patients included in the multicenter TMA study.** (A) Kaplan-Meier plot representing the overall survival according to substudy. (B) Kaplan-Meier plot representing the overall survival according to the adjuvant therapy received.

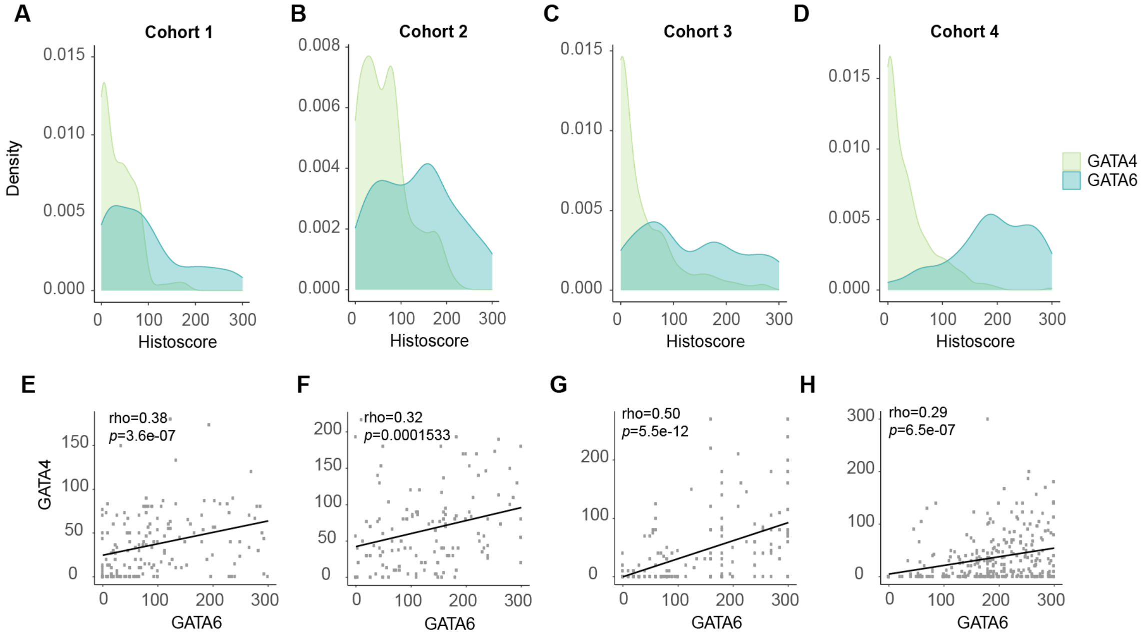


**Supplementary Figure 4. GATA4 and GATA6 distribution and correlation by substudy.** (A-D) Density plot depicting GATA4 and GATA6 histoscore distribution by substudy. (E-F) Scatter plot representing the correlation between GATA4 and GATA6 histoscores by substudy (Spearman correlation).

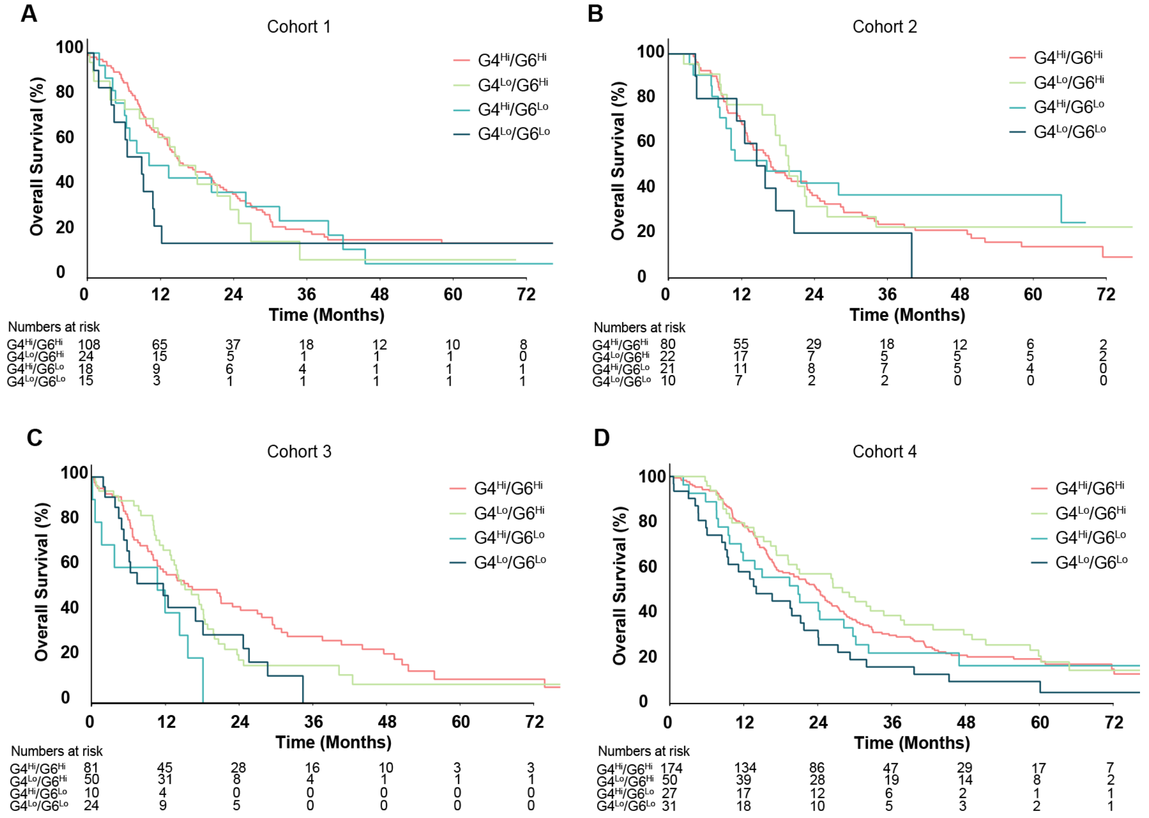


**Supplementary Figure 5. Patient survival according to GATA4 and GATA6 expression levels by substudy.** (A-D) Kaplan Meier estimates of overall survival by combined GATA4 and GATA6 IHC scores according to patient cohort.

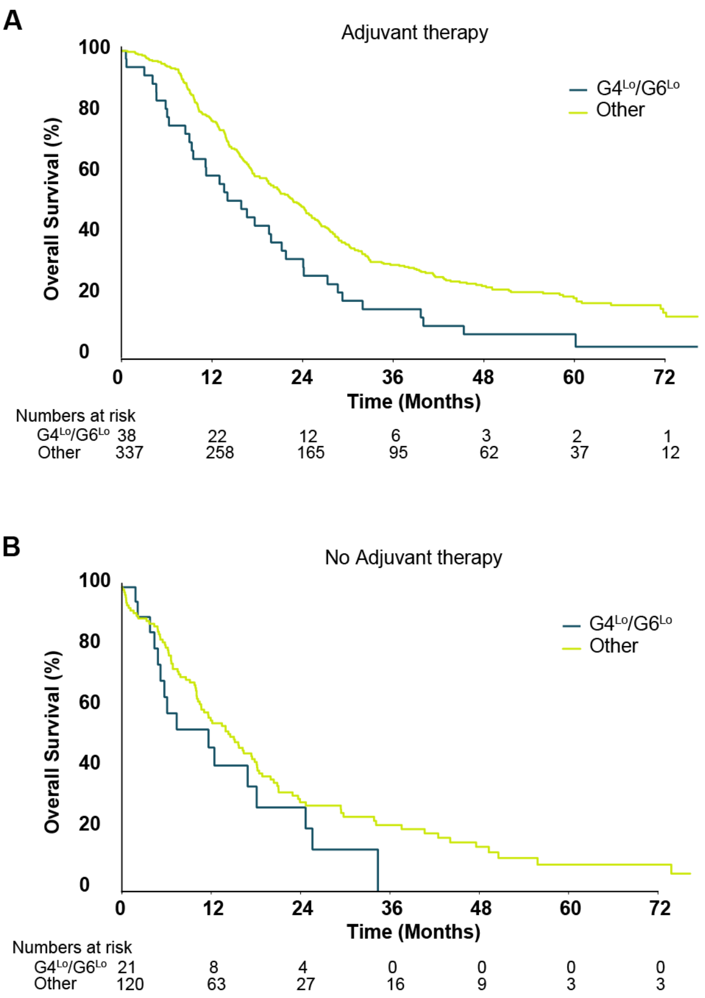


**Supplementary Figure 6. Survival analysis of patients according to adjuvant treatment and GATA4 and GATA6 expression levels.** (A,B) Kaplan-Meier plot representing the overall survival of patients with G4$^{Lo}$/G6$^{Lo}$ tumors vs. all others stratified according to whether they received adjuvant therapy or not.

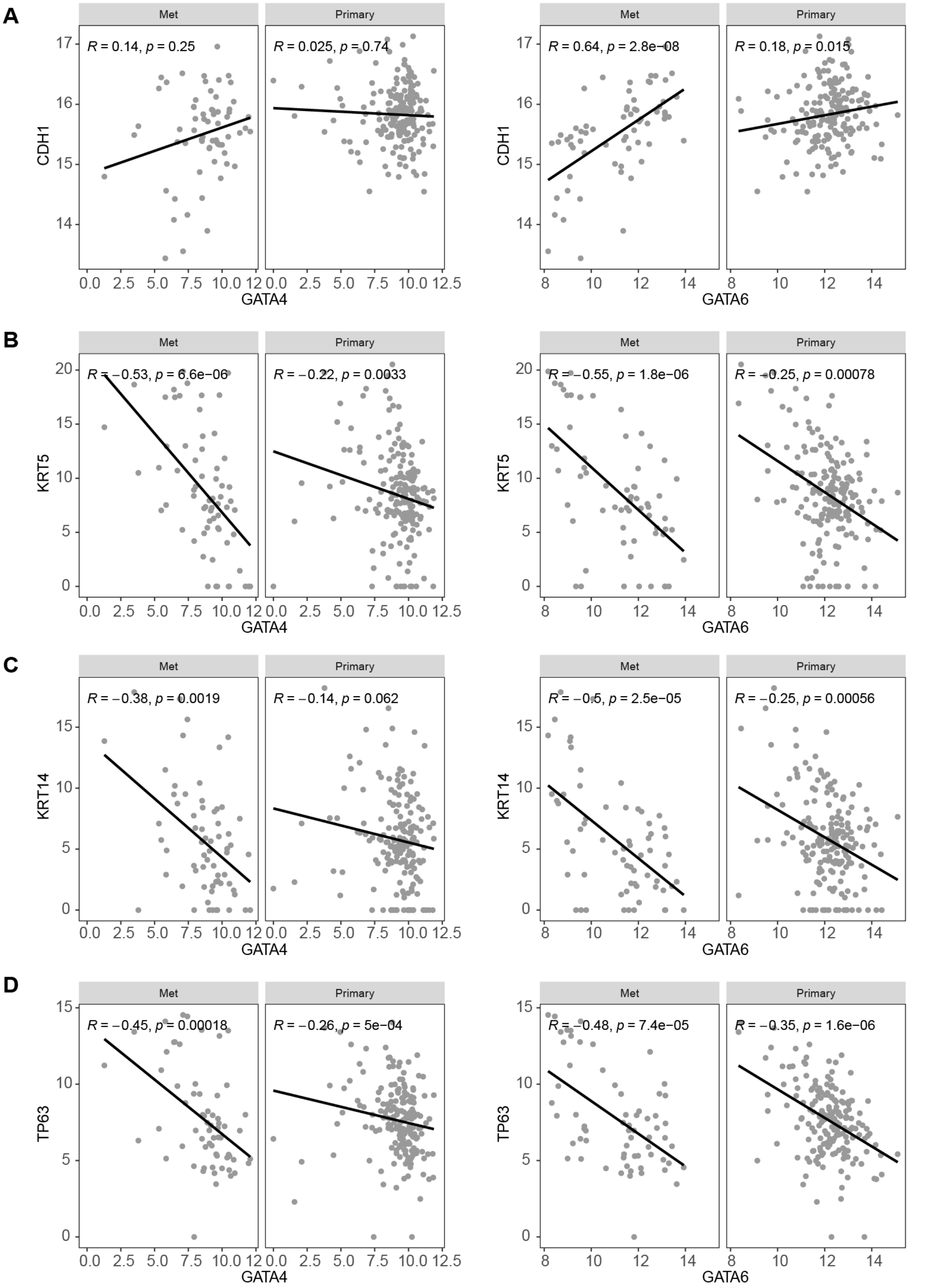


**Supplementary Figure 7. GATA4 and GATA6 expression correlates with basal genes in PDAC metastases.** (A-D) Scatter plot showing the correlation between GATA4 or GATA6 mRNA levels and classical (CDH1) or basal (KRT5/6, KRT14, TP63) genes in primary versus metastatic tumours (Spearman correlation).

#### SUPPLEMENTARY TABLES

**Supplementary Table 1. Patient cohorts included in the study.**

**Supplementary Table 2. Baseline demographics, clinical-pathological characteristics and expression of GATA4 and GATA6 in tumours from patients included in the study.**

**Supplementary Table 3. WGCNA analysis: individual genes included in the 33 modules identified (separate Excel sheet).**

**Supplementary Table 4. WGCNA analysis: functional annotation of the modules identified (separate Excel sheet).**

**Supplementary Table 5. Multivariable modeling of overall survival in the patients included in the study considering administration of adjuvant therapy.**

**Supplementary Table 1. Patient cohorts included in the study.**

| | Cores/tumour | Total number of cases | Number of cases included in the analysis* |
|---|---|---|---|
| Cohort 1 (Berlin) | 3 | 210 | 165 |
| Cohort 2 (Munich) | 3 | 162 | 133 |
| Cohort 3 (Regensburg) | 2 | 461 | 165 |
| Cohort 4 (ESPAC-3) | 4 | 434 | 282 |

* Only cases for which information on both markers was available were included in the analysis.

**Supplementary Table 2. Baseline demographics, clinical-pathological characteristics and expression of GATA4 and GATA6 in tumours from patients included in the study.**

| | | Summary statistics | | | | |
|---|---|---|---|---|---|---|
| VARIABLE | | Cohort 1 | Cohort 2 | Cohort 3 | Cohort 4 | TOTAL |
| Total | | 165 | 133 | 165 | 282 | 745 |
| Age | <65 | 86 (52%) | - | 85 (52%) | 160 (57%) | 331 (44%) |
| | >65 | 78 (47%) | - | 80 (48%) | 122 (43%) | 280 (38%) |
| | median (IQR) | 64.5 (58.7, 71.5) | - | 65 (57, 70) | 64 (57, 70) | 64 (57, 70.614) |
| Sex | Female | 76 (46%) | - | 72 (44%) | 121 (43%) | 269 (36%) |
| | Male | 89 (54%) | - | 93 (56%) | 161 (57%) | 343 (46%) |
| Stage | 1 | 0 (0%) | 2 (2%) | 6 (4%) | 17 (6%) | 25 (3%) |
| | 2 | 34 (21%) | 10 (8%) | 27 (16%) | 64 (23%) | 135 (18%) |
| | 3 | 121 (73%) | 101 (76%) | 123 (75%) | 191 (68%) | 536 (72%) |
| | 4 | 8 (5%) | 16 (12%) | 8 (5%) | 7 (2%) | 39 (5%) |
| Grade | 1 | 7 (4%) | 9 (7%) | 9 (5%) | 19 (7%) | 44 (6%) |
| | 2 | 89 (54%) | 60 (45%) | 53 (32%) | 185 (66%) | 387 (52%) |
| | 3 | 67 (41%) | 60 (45%) | 94 (57%) | 72 (26%) | 293 (39%) |
| Lymph Nodes | Neg | 38 (23%) | 39 (29%) | 50 (30%) | 54 (19%) | 181 (24%) |
| | Pos | 125 (76%) | 90 (68%) | 112 (68%) | 228 (81%) | 555 (74%) |
| Resec. Margins | R0 | 103 (62%) | 0 (0%) | 131 (79%) | 154 (55%) | 388 (52%) |
| | R1 | 42 (25%) | 0 (0%) | 30 (18%) | 128 (45%) | 200 (27%) |
| Adj. Therapy | Adj | 0 (0%) | 62 (47%) | 31 (19%) | 282 (100%) | 375 (50%) |
| | No Adj | 0 (0%) | 7 (5%) | 134 (81%) | 0 (0%) | 141 (19%) |
| GATA 6 | <150 | 110 (67%) | 68 (51%) | 75 (45%) | 69 (24%) | 322 (43%) |
| | >150 | 37 (22%) | 60 (45%) | 76 (46%) | 207 (73%) | 380 (51%) |
| | median (IQR) | 76.7 (26.7, 133.3) | 145 (60, 195) | 100 (60, 200) | 190.7 (146.8, 251.5) | 156.7 (66.7, 223.3) |
| GATA 4 | <30 | 45 (27%) | 30 (23%) | 29 (18%) | 90 (32%) | 194 (26%) |
| | >30 | 81 (49%) | 91 (68%) | 62 (38%) | 111 (39%) | 345 (46%) |
| | median (IQR) | 30 (3.3, 60) | 56.7 (30, 90) | 10 (0, 60) | 21.25 (0, 50) | 30 (0, 70) |

**Supplementary Table 5. Multivariable modeling of overall survival in the patients included in the study considering administration of adjuvant therapy.**

| Covariate | Level | HR (95% CI) | P-value |
|---|---|---|---|
| Grade | Poor | | |
| | Moderate | 0.73 (0.093, 3.314) | <0.001 |
| | Well | 0.81 (0.181, 1.104) | 0.269 |
| Stage | 1 | | |
| | 2 | 1.3 (0.293, 0.906) | 0.365 |
| | 3 | 1.39 (0.286, 1.155) | 0.248 |
| | 4 | 2.36 (0.337, 2.553) | 0.011 |
| Nodes | Negative | | |
| | Positive | 1.63 (0.112, 4.426) | <0.001 |
| Adjuvant therapy | Yes | | |
| | No | 1.50 (0.376, 1.079) | 0.281 |
| | Unknown | 2.02 (0.368, 1.919) | 0.055 |
| Resection margins | No | | |
| | Yes | 1.34 (0.104, 2.795) | 0.005 |
| | Unknown | 0.87 (0.265, 0.516) | 0.606 |
| Adjuvant therapy YES GATA 4/6 | Other | | |
| | Lo-Lo | 1.543 (5.128, 0.451) | 0.027 |
| Adjuvant therapy NO GATA 4/6 | Other | | |
| | Lo-Lo | 1.447(3.497, 0.774) | 0.196 |
| Adjuvant therapy Unknown GATA 4/6 | Other | | |
| | Lo-Lo | 1.919(3.559, 0.432) | 0.021 |
